# Fundamental Limits of Inferring Dynamical Gene Regulatory Models from Single-Cell Data

**DOI:** 10.1101/2025.09.12.674509

**Authors:** Ali Balubaid, Juan P. Bernal Tamyo, Rohit Kumar, Hafeez Agboola, David Gomez Cabrero, Narsis Kiani, Jesper Tegner

**Affiliations:** Biological and Environmental Science and Engineering Division, King Abdullah University of Science and Technology (KAUST), Thuwal 23955-6900, Saudi Arabia; Computer, Electrical and Mathematical Sciences and Engineering Division, King Abdullah University of Science and Technology (KAUST), Thuwal 23955-6900, Saudi Arabia; Algorithmic Dynamics Lab, Center of Molecular Medicine, Karolinska Institute, Stockholm, Sweden; Department of Oncology-Pathology, Karolinska Institutet, Stockholm, Sweden; Unit of Computational Medicine, Department of Medicine, Center for Molecular Medicine, Karolinska Institutet, Karolinska University Hospital, L8:05, SE-171 76, Stockholm, Sweden; Science for Life Laboratory, Tomtebodavägen 23A, SE-17165, Solna, Sweden

**Keywords:** Single-cell dynamics, RNA velocity, Gene regulatory networks, Jacobian estimation

## Abstract

A long-standing goal in systems biology is to construct computational models that capture gene regulatory dynamics. On the one hand, graph-based regulatory network models rely on prior knowledge but are constrained by unknown kinetic parameters. On the other hand, recent advancements in single-cell sequencing technologies, particularly RNA velocity and pseudotime analysis, offer data-driven approaches to infer kinetic parameters. However, accurate modeling of biological dynamics requires the estimation of the Jacobian matrix, which captures signal propagation within a regulatory system.

Several computational methods have recently emerged to infer Jacobians from single-cell data, including SpliceJAC, scMomentum, and Dynamo. Here, we evaluate these methods by analyzing Jacobian eigenvalues, assessing their ability to capture stability and oscillatory properties. Using simulated ground truth data, we find that methods such as Dynamo and scMomentum generate models that capture the behavioral properties of the cells, however, they do not reliably capture the structural properties.

Our results highlight the limitations of dynamic GRN inference, namely, the inability of kinetics to alleviate the need for further structural constrains for the models to adequately reflect the structural properties of the GRN. Developing more robust approaches for Jacobian estimation from single-cell data will be essential for building accurate, large-scale dynamical models of gene regulation.

**Graphical Abstract:** Overview of Benchmarking Jacobian-based inference in single-cell RNA Data.

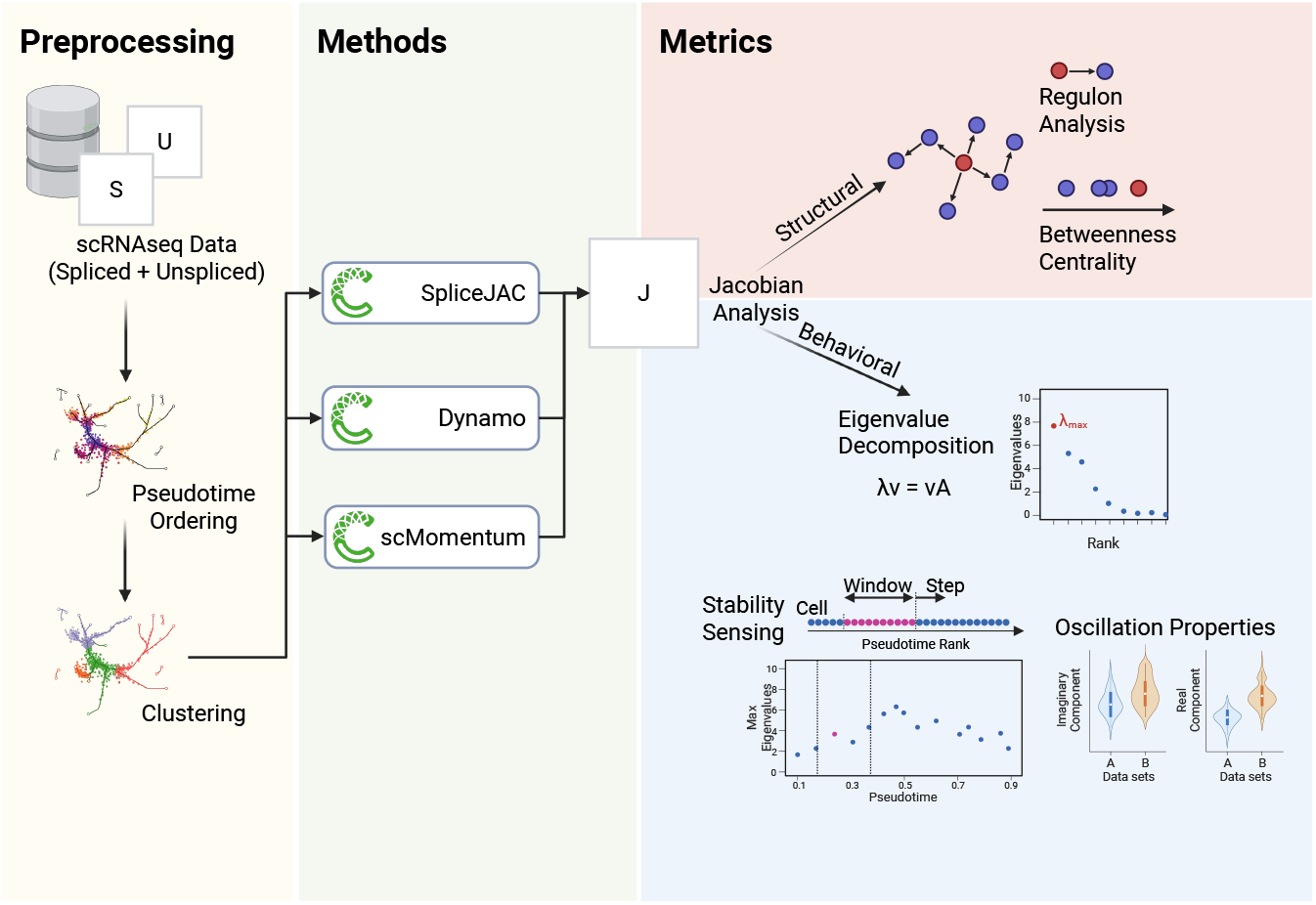

## Introduction

Single-cell RNA sequencing (scRNA-seq) has transformed our understanding of cellular heterogeneity and gene regulation by capturing high-dimensional transcriptomic states at single-cell resolution[1]. These datasets enable the reconstruction of cellular trajectories and the inference of gene regulatory networks (GRNs) but remain inherently static snapshots, limiting their ability to capture cellular dynamics fully [2, 3, 4].

RNA velocity introduced a significant advancement by inferring directional cell-state transitions based on spliced and unspliced mRNA ratios, providing a probabilistic framework for short-term fate predictions [5, 6]. However, while RNA velocity enhances trajectory inference, it lacks a mechanistic basis, as it does not explicitly model regulatory interactions governing these transitions[7]. In parallel, GRN inference approaches extract regulatory dependencies from single-cell data but remain fundamentally static, failing to describe how perturbations propagate through the network over time[3].

To bridge the gap between cellular transitions and underlying regulatory dynamics, Jacobian-based approaches have emerged as a potential solution[8]. The Jacobian matrix, a fundamental tool in dynamical systems theory, quantifies how small perturbations in gene expression propagate through regulatory interactions, capturing stability properties, oscillatory behaviors, and critical transitions [9]. Therefore, the Jacobian serves as a mechanistic bridge between RNA velocity and gene regulatory networks, providing a direct mapping of regulatory influences that drive cell fate transitions and lineage bifurcations[10]. Given its ability to describe system stability and state changes, the Jacobian can also predict critical transitions in cell differentiation[11].

Current computational frameworks, such as SpliceJAC, Dynamo, and ScMomentum, have been proposed to estimate Jacobians from scRNA-seq data, demonstrating their potential to recover gene regulatory interactions dynamically [12, 13, 14]. However, these methods have yet to undergo a systematic comparative analysis, leaving key questions unanswered: How accurately do they recover true regulatory interactions? How stable are their predictions across datasets? How robust are they to noise and parameter variations?

Here, we present a comprehensive benchmarking framework for Jacobian estimation methods in single-cell analysis. We systematically assess: (1) behavioral integrity, the inferred Jacobians’ capture of the system’s stability and oscillatory properties; (2) structural integrity, the inferred Jacobian’s reflection of the underlying GRN structure; and (3) computational robustness, evaluating sensitivity to parameter variations. By rigorously benchmarking these models, our study establishes critical guidelines for integrating single-cell transcriptomics with dynamical systems approaches, enabling more accurate inference of gene regulatory dynamics, highlighting current advantages and limitations for Jacobian-based approaches.

## Methods

### Data Generation and Preprocessing

#### MET Circuit Model

The mesenchymal-to-epithelial transition (MET) and its inverse (EMT) describe cell state transitions found in development and cancer[15]. We use the model shown in equation 1 as described by Barcenas[16] using the parameters from the original publication by Tian et al.[17], all listed in tables 1 and 2. In the model, each species is described using spliced and unspliced variables *U, V* where the generation component is used for the unspliced variable production, and the degradation rate is used for the degradation of the spliced variable.

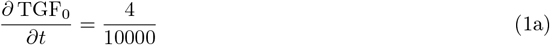

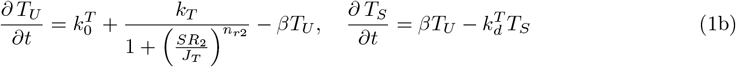

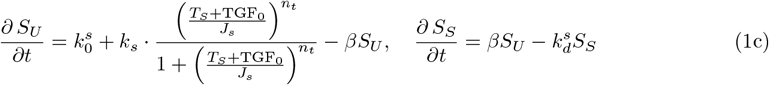

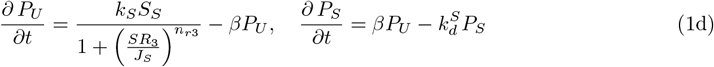

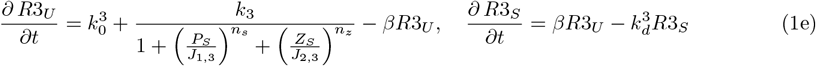

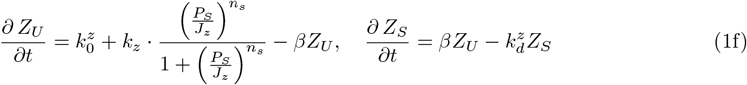

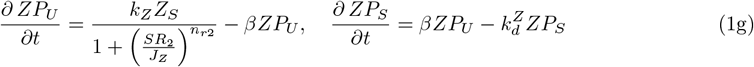

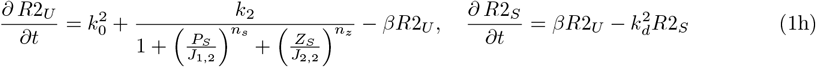

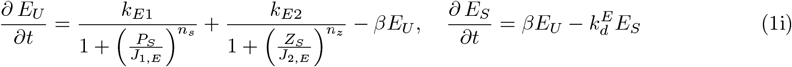

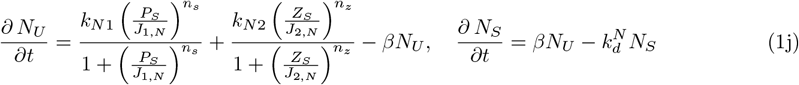

**Table 1:** Initial Conditions.

| Variable | Value | Variable | Value |
| --- | --- | --- | --- |
| $T_U(0)$ | 0.16 | $T_S(0)$ | 0.16 |
| $S_U(0)$ | 0.01 | $S_S(0)$ | 0.01 |
| $P_U(0)$ | 0.01 | $P_S(0)$ | 0.01 |
| $R3_U(0)$ | 0.38 | $R3_S(0)$ | 0.38 |
| $Z_U(0)$ | 0.03 | $Z_S(0)$ | 0.03 |
| $ZP_U(0)$ | 0.01 | $ZP_S(0)$ | 0.01 |
| $R2_U(0)$ | 0.35 | $R2_S(0)$ | 0.35 |
| $E_U(0)$ | 3.20 | $E_S(0)$ | 3.20 |
| $N_U(0)$ | 0.00 | $N_S(0)$ | 0.00 |

**Table 2:**
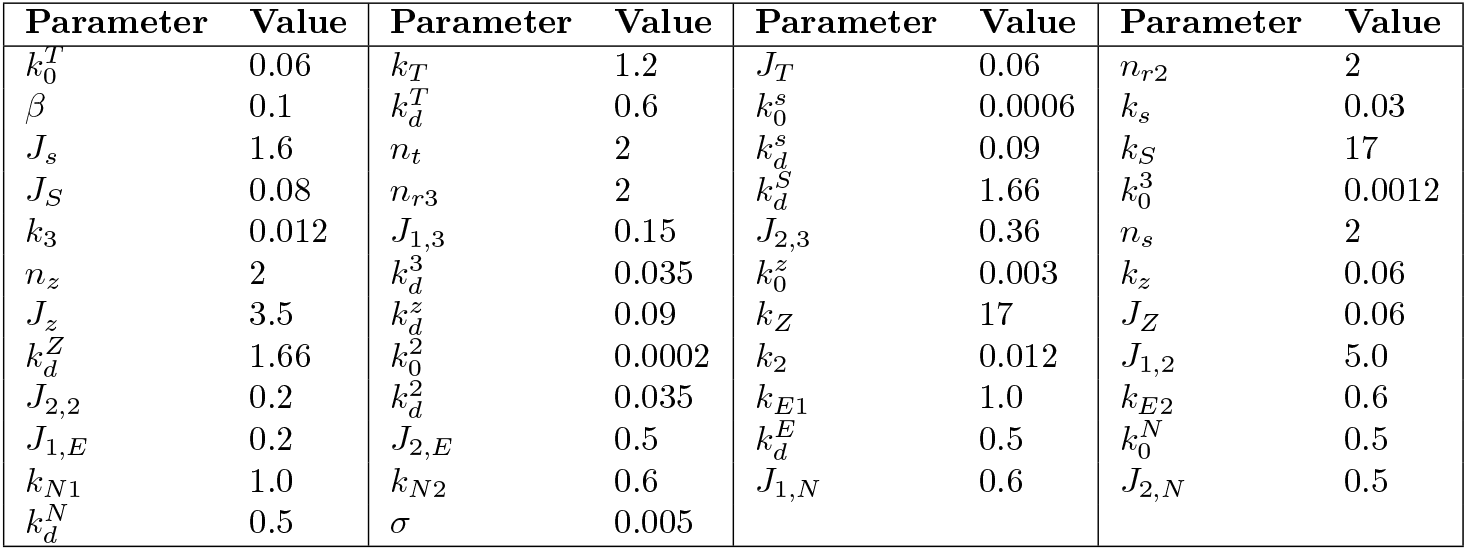
Model Parameters.

| Parameter | Value | Parameter | Value | Parameter | Value | Parameter | Value |
| --- | --- | --- | --- | --- | --- | --- | --- |
| $k_0^T$ | 0.06 | $k_T$ | 1.2 | $J_T$ | 0.06 | $n_{r2}$ | 2 |
| $\beta$ | 0.1 | $k_d^T$ | 0.6 | $k_0^s$ | 0.0006 | $k_s$ | 0.03 |
| $J_s$ | 1.6 | $n_t$ | 2 | $k_d^s$ | 0.09 | $k_S$ | 17 |
| $J_S$ | 0.08 | $n_{r3}$ | 2 | $k_d^S$ | 1.66 | $k_0^3$ | 0.0012 |
| $k_3$ | 0.012 | $J_{1,3}$ | 0.15 | $J_{2,3}$ | 0.36 | $n_s$ | 2 |
| $n_z$ | 2 | $k_d^3$ | 0.035 | $k_0^z$ | 0.003 | $k_z$ | 0.06 |
| $J_z$ | 3.5 | $k_d^z$ | 0.09 | $k_Z$ | 17 | $J_Z$ | 0.06 |
| $k_d^Z$ | 1.66 | $k_0^2$ | 0.0002 | $k_2$ | 0.012 | $J_{1,2}$ | 5.0 |
| $J_{2,2}$ | 0.2 | $k_d^2$ | 0.035 | $k_{E1}$ | 1.0 | $k_{E2}$ | 0.6 |
| $J_{1,E}$ | 0.2 | $J_{2,E}$ | 0.5 | $k_d^E$ | 0.5 | $k_0^N$ | 0.5 |
| $k_{N1}$ | 1.0 | $k_{N2}$ | 0.6 | $J_{1,N}$ | 0.6 | $J_{2,N}$ | 0.5 |
| $k_d^N$ | 0.5 | $\sigma$ | 0.005 | | | | |

The conversion between unspliced and spliced is modulated by parameter *γ* = 1 for all species. The splicing rate coefficient is assumed to be rescaled to the model value. The model bifurcates by varying exogenous TGF concentration (*TGF*_0_) by gradually increasing from 0 to 4. The stochastic model was simulated using the Euler-Maruyama algorithm.

We first generate data sets corresponding to a time series, sample the fast bifurcating process, and sample the slow bifurcating process. We produced 50 instances of each data form. We use the MET circuit model, following the schematic in Figure 1:

**Figure 1:**
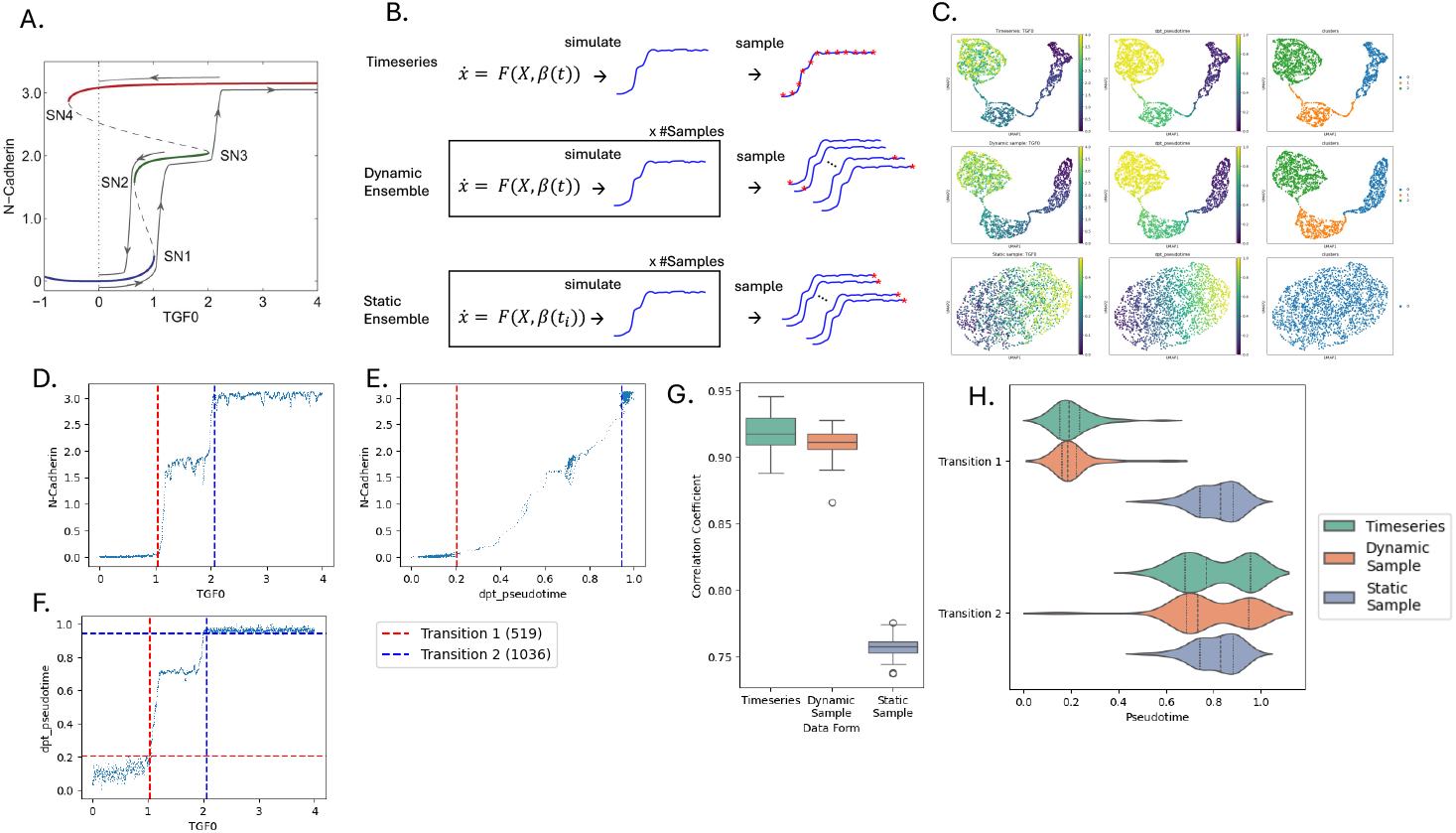
Validation of approximating the temporal evolution using Pseudotime: A. The phase plot shows stable states for the N-Cadherin state variable along the bifurcation parameter TGF0, revealing the three steady states colored blue, green, and red. B. Schematic showing the generation of the three data forms using the same model. C. UMAP of dynamic sampling colored by the bifurcation parameter TGF0, diffusion pseudotime, and identified clusters for the three models. D. State variable N-cadherin versus bifurcation parameter TGF0 while identifying transition points 1 and 2 with red and blue lines, respectively. E. State variable N-cadherin versus diffusion pseudotime while identifying transition points 1 and 2 with red and blue lines, respectively. F. Diffusion pseudotime versus bifurcation parameter TGF0 while identifying transition points 1 and 2 with red and blue lines for both axes, respectively. G. Assess the accuracy of pseudotime using the correlation between the bifurcation parameter TGF0 and pseudotime. H. Transition pseudotime distribution for the different data forms.

- **Time series data:** Time-series data are temporally dependent and, therefore, are generated from a single simulation. The bifurcation parameter has a constant rate of change in this system. The data were then down-sampled to 2000 samples to evaluate downstream. *X* = *F* (*x*) such that 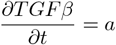 where a (default *a* = 0.004) is a constant rate of change of bifurcation variable *TFGβ*.
- **Sampled Dynamic (Fast-bifurcating) process:** Unlike time series, the data points are temporally independent but originate from the same process. Therefore, we run 2000 simulations and sample each at a given time interval to replicate real-world single-cell data. The bifurcation parameter has a constant rate of change in this system. *X F* (*x*) such that 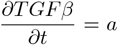 The values are the same as eq 1.
- **Sampled Static (Slow-bifurcating) process:** Similar to the fast-bifurcating process, we run 2000 simulations. However, the bifurcation parameter has a rate of change of zero in this system; rather, the bifurcation parameter is set at interval values along the bifurcation axis for each simulation. The system is allowed to converge, and the terminal steady state is sampled. *X F* (*x*|*b*) such that 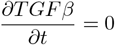

#### Preprocessing

We process each generated data table by a standard single-cell pipeline using scanpy (v1.10.2)[18]. The data is filtered, normalized, and leiden[19] clustered. We also infer a pseudotemporal ordering using diffusion pseudotime[20], and visualize in a UMAP embedding[21].

#### DynGen In Silico Simulation[22]

To compare oscillation detection and GRN inference ability, we generate two *in silico* datasets using the DynGen package (v 1.0.5)[22] for the cycle and the linear backbones composed of 30 TFs, 100 target genes, and 100 housekeeping genes. A dataset with 2000 cells is generated with default configuration parameters. The datasets were normalized, scaled, dimensionally reduced, and clustered using the scanpy package (v1.10.2)[18], the cell with the lowest simulation time was assigned as root, and diffusion pseudotime was computed. The Jacobian inference methods were then applied, with the full dataset treated as a cluster.

To assess the methods’ ability for oscillatory behavior modeling, a series of metrics are computed, including the eigenvalue magnitude and phase distributions to evaluate the Jacobian dynamic range, imaginary and real component distributions to evaluate oscillatory and scaling behaviors, and a fraction of negative real component distribution to determine stability.

For the regulon evaluation, we select the edges with the top weights. A regulon comprises a regulator (i.e., TFs) regulating an effector (i.e., Target gene). In our matrix with 230 genes, we select the top 1000 regulons by weight for the ground truth network and the Jacobian matrix estimated by the different methods. The sets of top regulons are compared using the Jaccard similarity index for the regulator, the effector, and the combination of both (i.e., the regulon).

Next, the network inference is evaluated by comparing the inferred Jacobians to the known ground truth. The quality of inference is assessed by using a subset of metrics used for GRN benchmarking[3], estimation of the area under the receiver operator curve (AUROC), the area under the precision-recall curve (AUPR), and the early precision-recall (EPR) for edge detection. A more detailed description of the different metrics can be found in the original publication [3]. EPR was computed using the top-k edges, which in our case is defined as the number of non-zero edges in the real network.

#### Spermiogenesis Differentiation[23]

A spermatogenic wild-type differentiation data was used since it provides a linear progression along differentiation. In our analysis, we use the boundaries to reference cell-type transitions and evaluate the metric trends in reference. The data was QCed (keep cells >700 total counts, >200 features, keep features in >20 cells) and preprocessed with the monocle recipe preprocessing. The data was then annotated manually using markers from source publications. For a linear progression, isolated clusters were removed, and diffusion pseudotime was computed using the scanpy package (v1.10.2)[18]. Velocity values were then computed for the data for downstream analysis[6]. Jacobian inference methods were applied, and the trends were evaluated with respect to the transition between cell type distributions. The dataset was also used for sensitivity analysis of gene selection, which we describe in a later section.

#### Mouse Pancreatic Endocrinogenesis[24]

The mouse pancreatic endocrinogenesis dataset describes the emergence of Alpha, Beta, Delta, and Epsilon cell types from the Ductal cells[24]. The dataset has been commonly leveraged for RNA velocity application[6, 14]. The data used was acquired from the ScVelo (v0.3.3)[6] sample dataset repository with 3696 cells and 27998 features, then preprocessed and velocity model parameters computed. The diffusion pseudotime was computed using scanpy (v1.10.3)[25]. The data was then analyzed using the different Jacobian inference methods.

### Benchmarked Tools

#### SpliceJAC[13]

SpliceJAC leverages the collective dynamics of spliced and unspliced RNA to infer the Jacobian using a splicing model, shown in eqs. (2a) and (2b):

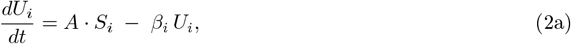

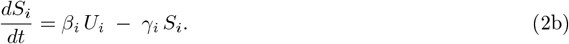

By assuming the rate of change of unspliced RNA is zero 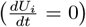, the estimation of *A* becomes a regression problem (e.g., linear, ridge, or lasso). Additionally, SpliceJAC assumes the splice conversion rate coefficient is identical for all species, so *β*_*i*_ = *β* = 1 (via time rescaling). This enables direct computation of 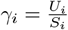.

Using these assumptions, SpliceJAC assembles the Jacobian for spliced and unspliced variables for each gene, as indicated in eq. (2), and the Jacobian is inferred independently for each cell type.

#### Dynamo[14]

Dynamo fits a velocity vector function *v* = *f* (*x*) using a summed basis vector kernel based on the velocity data. This results in a differentiable geometric space, upon which regulatory elements 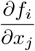 can be inferred. The components are assembled to realize a Jacobian for the states of interest. By assuming that the dynamics occur on a manifold, Dynamo predicts trajectory paths and perturbation shifts. Therefore, given the state of a system and the vector function, a Jacobian could be estimated for each cell.

#### ScMomentum[12]

ScMomentum uses the Hopfield network to fit a model to the RNA velocity data. The velocity is computed from spliced and unspliced count matrices using a velocity quantification method to estimate other splicing coefficients (i.e., estimate gamma, beta, etc.). The Hopfield model then fits the data, inferring the relationships between the different genes. We modify the code to compute the Jacobian by using the derivative of the activation function. Jacobians for individual cells are computed by taking the model’s derivative using the cell’s state and the cell-type-specific model.

### Sensitivity Analysis

#### Window Sensitivity

One potentially impacting hyperparameter is the window size. We, therefore, test a range of window sizes for the different methods and how they affect the profile of the leading eigenvalues over pseudotime. While for some methods, the window size is only implemented after the computation of the eigenvalues for averaging (i.e., ScMomentum, dynamo), others use the window to estimate the Jacobian (i.e., SpliceJAC) and would therefore be expected to have more influence. A sampled dynamic dataset was used for the analysis with a fixed step size (step size = 50) and varying window sizes (window sizes = 50, 75, 100, 150, 200, 250, 300). The leading eigenvalue profile was reported for each method separately.

#### Step Sensitivity

Another potentially impacting hyperparameter is the step size. Similar to the previous analysis, a range of step sizes was tested for the different methods and how they impact the profile of the leading eigenvalues over pseudotime. The exact expectations hold for the step sizes as the window sizes, with the step size being implemented after the computation of the eigenvalues for averaging (i.e., ScMomentum, Dynamo). At the same time, it directly estimates the choice of cells for the Jacobian estimation (i.e., SpliceJAC) and is therefore expected to have more influence. The analysis was done in a sampled dynamic dataset with a fixed window size (window size = 100) and varying step sizes (step sizes = 10, 20, 30, 40, 50, 60, 70). The leading eigenvalue profile was reported for each method separately.

#### Cluster Sensitivity

The method ScMomentum initiates its Jacobian estimation by computing the Hopfield for every cluster. Downstream results could, therefore, be influenced by the clustering resolution. We test the clustering sensitivity by estimating the Jacobian for different clustering resolutions using the Louvain algorithm (resolution = 0.1, 0.3, 0.8, 1, 3, 5) and observing the leading eigenvalue profile given the number of clusters in the data.

#### Gene Number Sensitivity

To assess the impact of the number of genes on the different methods, we tested for a range of genes using the spermiogenesis dataset [23]. Some methods, such as SpliceJAC, rely on a small subset of genes and are tested on a lower range (ngenes=20, 30, 40, 60, 80), while others require more genes for computations and are tested on a higher range (ngenes=100, 200, 300, 400, 500).

#### Cell Sampling Sensitivity

We assess the impact of the number of cells on the resolution of the trends using the data generated with the MET in silico model. Jacobian inference methods were run for 500, 2000, 5000, and 10000 cells.

## Results

### Pseudotemporally Ordered Dynamic Sampled Cells Approximate the Time Series

To systematically assess the ability of Jacobian-based methods to detect critical transitions from single-cell data, we implement an in silico model of the Mesenchymal-Epithelial Transition (MET) (see Methods) [17, 16]. This model describes progression through three distinct cell states, mediated by two-fold bifurcations. During each bifurcation, a stable and an unstable steady state converge and vanish in a saddle-node bifurcation, leading to a shift in cell state. The transitions are driven by external levels of transforming growth factor (TGF0), which serves as the bifurcation parameter of the system (Figure 1A). This model enables the simulation of three distinct data forms: time series data tracking a bifurcating system along a time-dependent parameter, dynamic sampling at discrete time points, and static sampling at steady states along the bifurcation axis (Figure 1B-C). This setup allows us to assess the limitations of pseudotemporal ordering in recovering trends similar to time series.

We first identify the correspondence between the known transition driver (TGF0), the computed pseudotime, and their impact on the system transition represented by changes in the state-variable N-Cadherin (similar to what is shown in Figure 1A). In our initial simulations under various starting conditions, we identified critical TGF*β* transition values at 1.04 and 2.10, corresponding to saddle-node bifurcations, the loss of a stable state at the gain of another, with a high correspondence between external TGF levels and pseudotime values (Figure 1D-E). We observe a loss of correspondence between pseudotime and the bifurcation parameter during steady state, in which variability is better attributed to the inert heterogeneity of the system than to a difference between states (Figure 1F). Nonetheless, the correlation between the bifurcation parameter values and pseudotime shows that static samples have a reduced ordering accuracy compared to time series and dynamic sampling approaches (Figure 1G). Additionally, we observe that the distribution of pseudotime around the first transition and second transitions is similar for time-series data and dynamically sampled data. In contrast, static data have cells with the same pseudotime values from both the first transition (cells with TGF0∼ 1.04) and the second transition (cells with TGF0∼ 2.10)(Figure 1H). We also observe a bimodal distribution for pseudotime values around transition 2, with modes occurring at 0.7 and 0.95 dimensionless pseudotime units (Figure 1H). This is likely the result of the noisy bistable system, where one mode corresponds to cells remaining in the intermediate steady state and others crossing to the destination steady state, leading to two possible pseudotime assignments for cells with the same TGF0 levels.

These findings suggest that pseudotemporal ordering can reliably recover the correct sequence of states when transitions occur faster than changes in the underlying driver. In such cases, the system may reasonably be treated as a time series, enabling a practical Jacobian estimation in downstream analyses. Having outlined the structural differences among static, dynamic, and time-series data, we are now in a position to assess how these distinctions influence the performance of various Jacobian estimation methods in detecting critical transitions.

### Expression Dynamic Range Determines Detectability of Transition Indicators

Transitioning out of steady cellular states is a fundamental process in multicellular systems [26]. Such transitions are effectively characterized by an increase in the real part of the leading eigenvalue of the Jacobian matrix, indicating the loss of stability in the modeled system [27]. Here, we assess the capacity of existing Jacobian inference methods, SpliceJAC, ScMomentum, and Dynamo, to detect these transition events within scRNA-seq data[13, 12, 14]. Transition detection performance differed markedly across Jacobian estimation methods. The analytical ground truth Jacobian reliably identified the second transition within the time series data but exhibited limited sensitivity to the first transition (Figure2A). This establishes an upper bound, indicating that even the true Jacobian fails to capture the first transition. Consequently, method evaluation focuses primarily on their ability to detect the second transition. The leading eigenvalue of SpliceJAC’s Jacobian fails to show clear peaks corresponding to the second transition (Figure 2B). In contrast, the leading Jacobian eigenvalues estimated using Dynamo and ScMomentum successfully capture the second transition using both time series and dynamic ensembles data forms, though with slightly different temporal positioning compared to the analytical method (Figure 2C, D). Our results indicate that ScMomentum and Dynamo are more likely to detect transition events, and therefore, we prioritize their results when analyzing published data.

**Figure 2:**
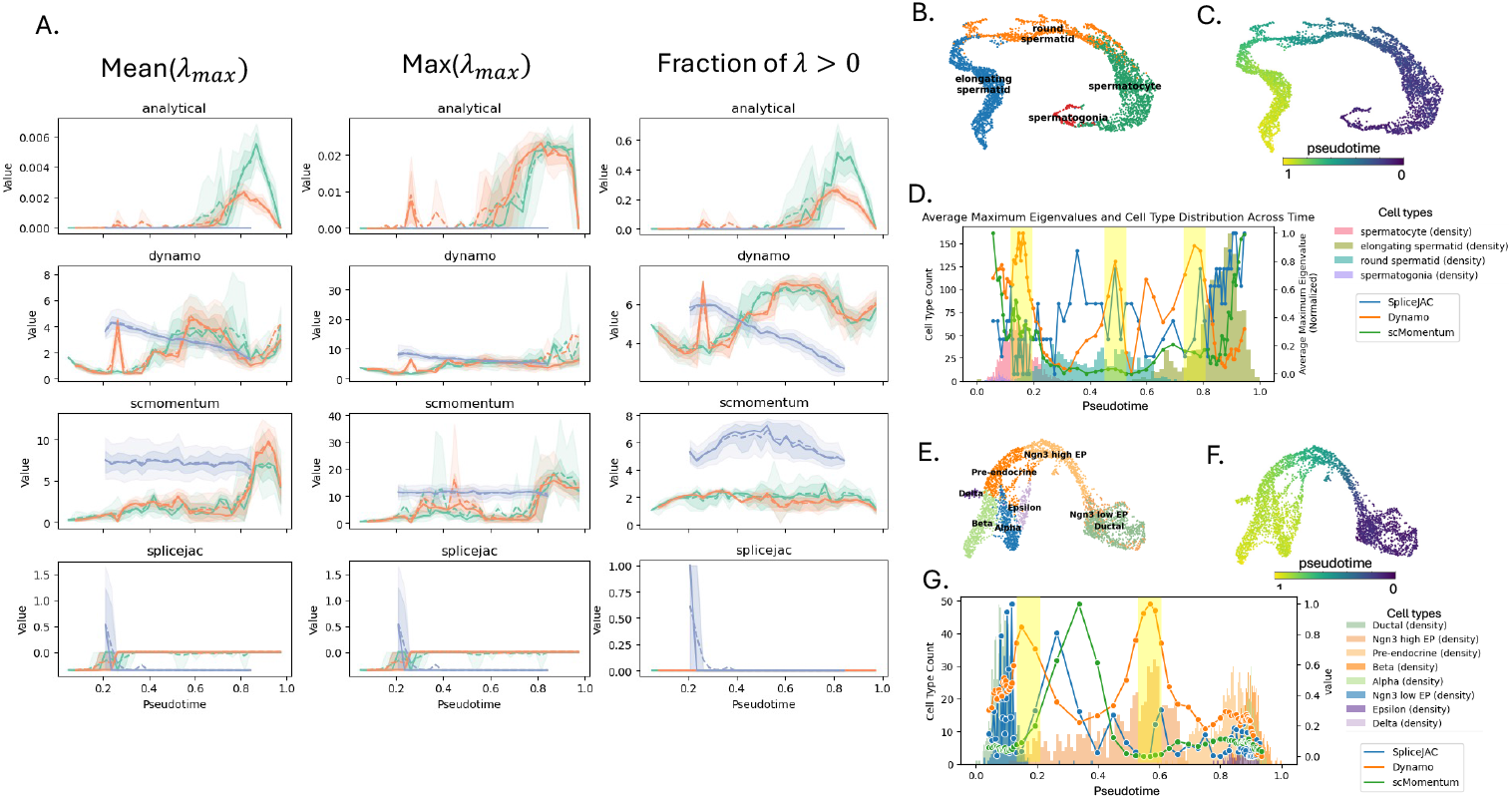
Behavioral Integrity - Transitions: **A**. Trends of the eigenvalue-based metrics derived from computed Jacobians across 50 datasets of identical form. Each line represents a specific data format: Timeseries, dynamic sampling, or static sampling. Columns show the metrics (from left to right): the average leading eigenvalue, the maximum leading eigenvalue, and the fraction of positive eigenvalues. Rows (from top to bottom) correspond to different Jacobian estimation methods: Analytical (control), Dynamo, ScMomentum, and SpliceJAC. **B, C**. UMAP projection of Spermiogenesis data colored by dataset-specific cell clusters (**B**) and annotated cell type labels (**C**). **D**. Leading eigenvalues over pseudotime plotted against cell type density for Spermiogenesis data. Yellow highlights indicate expected transition points. **E, F**. UMAP projection of mouse pancreatic endocrinogenesis data colored by cell type labels (**E**) and pseudotime values (**F**). **G**. Leading eigenvalues along pseudotime overlaid on cell type density for the same dataset, with expected transition points marked in yellow.

The analytical Jacobian implemented in time series data as the reference, all metrics show low error, except for the largest *λ >* 0. Instead, the metric of average *λ*_*max*_ shows the best results with the analytical Jacobian. ScMomentum shows the best results compared to the positive control, however, the performance seems particular to the use of the largest *λ*_*max*_ across pseudotime. Dynamo, on the other hand, shows more consistent performance across the used metrics.

Next, we infer the Jacobian along the spermatogenesis trajectory in mice[23], which involves the sequential progression from spermatogonia to spermatocytes, followed by round spermatids and ultimately elongating spermatids (Figure 2H–I). The Jacobians’ leading eigenvalues are detected for multiple methods at inter- and intra-cluster boundaries. While eigenvalue peaks between clusters are expected, large peaks describe a rapid early transition (as ordered by pseudotime) of spermatogonia to spermatocytes to round spermatids, suggesting lower state stability of spermatocytes. Indeed, the progression from spermatogonia to spermatid includes transit-amplifying meiotic and mitotic divisions, which could explain the overall loss of stability for the intraclusteral transition[28]. Another prominent peak in the Jacobian eigenvalue landscape is between round and elongating spermatids, indicating three separate steady states (Figure 2J). These correspond with previously reported distinct phenotypes of early round, late round, and elongating spermatids[29].

In pancreatic endocrinogenesis[24], Ductal cells give rise to endocrine progenitors (EP) with low Ngn3 expression, which then increase Ngn3 expression before transitioning to pre-endocrine cell type. Pre-endocrine cells eventually give rise to alpha, beta, delta, and epsilon cell types (Figure 2J-K). Dynamo and SpliceJAC reveal a peak between Ngn3 low and Ngn3 high PE cells, between Ngn3 high PEs and pre-endocrine cells, and lower peaks from PE cells to the terminal endocrine cell types (i.e., alpha, beta, delta, epsilon). This agrees with previous expectations, highlighting the boundaries between cell types. ScMomentum, on the other. hand peaks at an intermediate state in Ngn3 high PE cells, contradicting the other two methods (Figure 2L).

Our results demonstrate successful transition detection using ScMomentum and Dynamo Jacobian, given the analytical Jacobian baseline, whereas SpliceJAC exhibits inconsistent performance. Furthermore, the methods were applied to published scRNA-seq datasets, revealing their utility for stability analysis of cellular differentiation trajectories. Next, we evaluate whether current Jacobian inference methods can capture other complex dynamic behaviors.

### Pre-estimated Vector Fields Increase Oscillatory Property Capture

Jacobian analysis provides insights into system behavior near steady states. Beyond detecting transitions through peaks in the leading eigenvalue, the full eigenvalue spectrum encodes richer information about underlying regulatory dynamics. Notably, eigenvalues with non-zero imaginary components are indicative of oscillatory behavior.

To assess whether current Jacobian inference methods can capture such dynamics, we generated simulated datasets from ground truth gene regulatory networks (GRNs) exhibiting either linear or cyclic topologies (Figure 3A) [22]. For each inferred Jacobian, we examined three properties: (1) the proportion of real eigenvalues below zero (reflecting local stability), (2) the distribution of real component values, and (3) the phase angle (i.e., the ratio of imaginary to real components), which signals dominant dynamical regimes. The magnitude of the eigenvalues was used to evaluate the overall strength of the dynamics.

**Figure 3:**
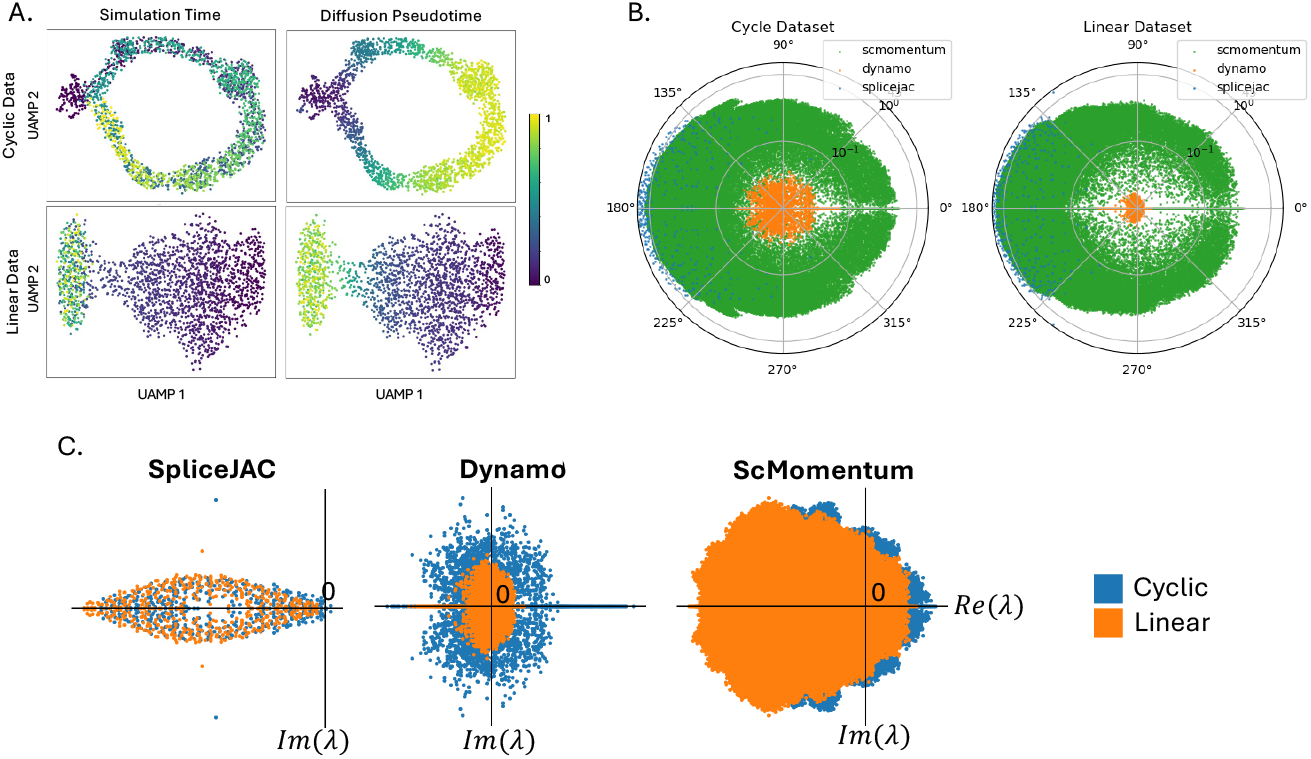
Behavioral Integrity - Oscillations: A. DynGen generated datasets projected on UMAP colored by simulation time (left) and diffusion pseudotime (right) for cyclic (top) and linear (bottom) backbones. C. Oscillatory property capture analysis for SpliceJAC for a fraction of negative real eigenvalues, eigenvalue phase distribution, eigenvalue magnitude distribution, eigenvalue real component distribution, and eigenvalue imaginary component distribution in that order from left to right. D, E.Similar to C for (D) Dynamo, and (E) ScMomentum.

Initial inspection of the eigenvalue distributions in phase space revealed marked differences between Jacobians inferred from linear and cyclic systems (Figure 3B). SpliceJAC and Dynamo exhibited broader spreads along the imaginary axis for cyclic trajectories, indicating their ability to capture oscillatory behavior. In contrast, ScMomentum showed minimal differentiation between linear and cyclic cases. Additionally, all methods displayed stronger positive real components under cyclic conditions, pointing to more destabilizing regulatory influences.

Across all methods, the fraction of negative real eigenvalues was consistently higher in linear systems, consistent with theoretical expectations (Figure 3C). For instance, SpliceJAC and ScMomentum showed distributions skewed further into the negative real domain for linear dynamics. These methods also exhibited more frequent non-zero real components, as confirmed by their eigenvalue magnitude distributions. To support this claim, we generate multiple dataset instances with cyclic and linear cell dynamics, and generate the eigenvalues for their inferred Jacobians. Clustering the datasets using the different properties of the eigenvalues shows that Dynamo, in particular, reliably captures the differences between the two dynamics (Table 3).

**Table 3:** Mann–Whitney U test comparing inter- vs intra-cluster distances between cyclic and linear datasets across Jacobian-derived metrics. Significance indicators: * *p <* 0.05, ** *p <* 0.01, *** *p <* 0.001.

| Method | Metric | Mean<br>(Between) | Mean<br>(Within) | U<br>Statistic | $p$-<br>value | Significance |
| --- | --- | --- | --- | --- | --- | --- |
| SpliceJAC | $\text{Re}(\lambda)$ | 6.6e-2 | 6.7e-2 | 2.6e5 | 4.1e-1 | |
| | $\text{Im}(\lambda)$ | 5.7e-2 | 5.8e-2 | 2.6e5 | 5.0e-1 | |
| | $\#(\text{Re}(\lambda) < 0)$ | 1.6e-2 | 1.7e-2 | 2.5e5 | 6.8e-1 | |
| Dynamo | $\text{Re}(\lambda)$ | 2.8e-4 | 2.5e-4 | 2.9e5 | 3.8e-7 | *** |
| | $\text{Im}(\lambda)$ | 8.7e-5 | 8.3e-5 | 2.8e5 | 3.9e-3 | ** |
| | $\#(\text{Re}(\lambda) < 0)$ | 1.7e-2 | 1.7e-2 | 2.5e5 | 6.2e-1 | |
| scMomentum | $\text{Re}(\lambda)$ | 1.4e-1 | 1.41e-1 | 2.6e5 | 4.7e-1 | |
| | $\text{Im}(\lambda)$ | 6.4e-2 | 6.5e-2 | 2.6e5 | 4.2e-1 | |
| | $\#(\text{Re}(\lambda) < 0)$ | 1.4e-2 | 1.3e-2 | 2.7e5 | 5.8e-2 | |

In terms of phase characteristics, SpliceJAC and ScMomentum displayed pronounced dynamical biases, while Dynamo presented a more uniform dynamic range (Figure 3C–E). Furthermore, the broader imaginary distributions observed in SpliceJAC and Dynamo for cyclic datasets suggest a stronger capacity to resolve complex, oscillatory dynamics, particularly those shaped by RNA velocity signals.

These findings highlight the advantages of RNA velocity-informed methods, especially Dynamo and SpliceJAC, in capturing cyclic regulatory behavior. This sensitivity is particularly important for modeling oscillatory gene expression programs, which are critical in developmental and homeostatic contexts.

Having evaluated transition and oscillatory behaviors, we next assess whether these Jacobian-based methods can accurately recover the underlying structure of gene regulatory networks.

### Direct Estimation of Jacobians Fails to Capture Regulatory Structure

We evaluated the structural integrity of the inferred Jacobians by comparing their recovered gene regulatory networks (GRNs) against ground truth data. This assessment focused on both regulon-level agreement and broader network-level properties.

An initial analysis of weight distributions revealed that inferred Jacobians typically had lower average interaction strengths compared to the ground truth and lacked similar distributional patterns (Figure 4A, S1A). Moreover, inferred networks tended to be denser, with SpliceJAC producing the sparsest matrices, a result of thresholding small-magnitude entries (Figure 4B, S1B).

**Figure 4:**
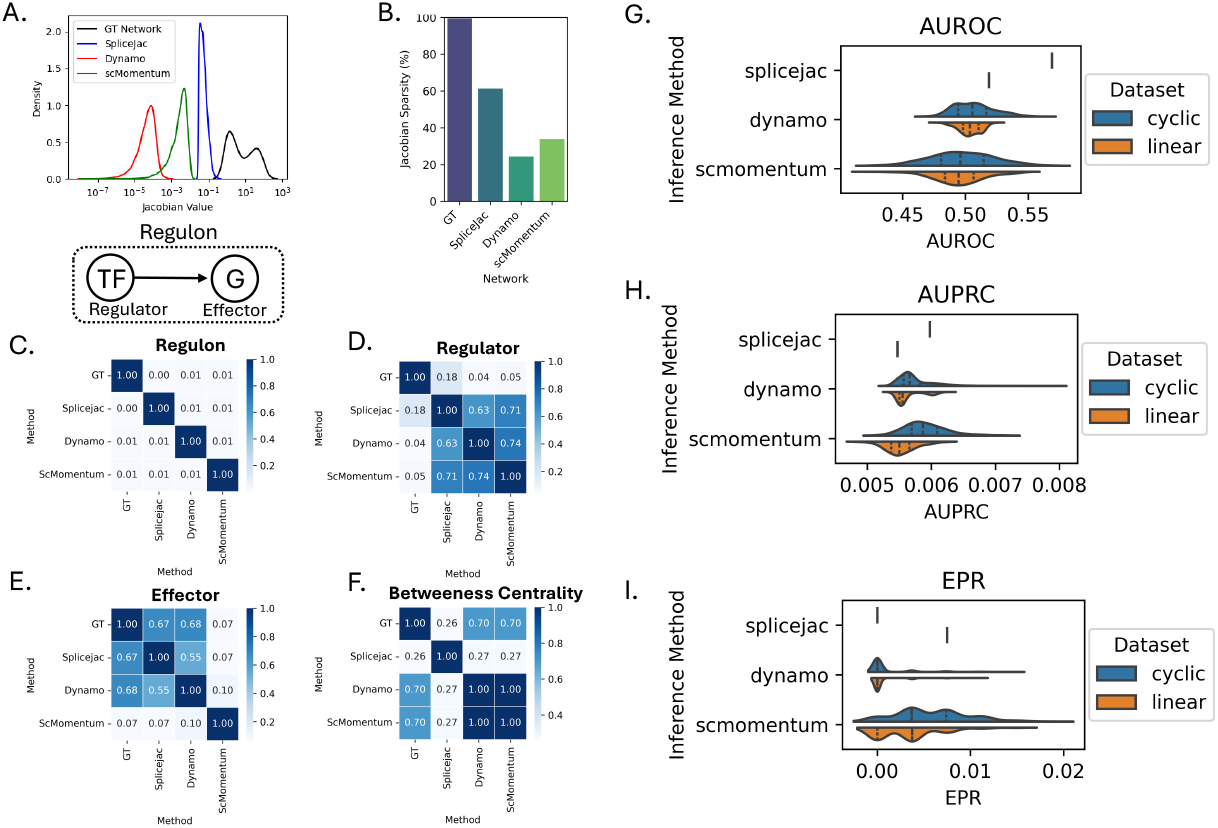
Structural Integrity: A. GRN weight distribution for the GT and Jacobians estimated by the other methods. B. The percentage of zeros in the Jacobians is estimated between the different methods compared to the ground truth. C. The network substructure similarity was evaluated for the top regulons (edge from regulator to effector), and the Jaccard index is computed between the top regulons, effectors, and regulators in order of edge value (Top_N_regulons = 1000). The similarity in the top genes by betweenness centrality was also evaluated (Top_N_genes = 115). D. The similarity of the overall Jacobian structure was quantified with AUROC, AUPRC, EPR, and EPR (activation regulons). For Dynamo and ScMomentum, the average Jacobian for all cells was used. All results correspond to the linear trajectory. GT: ground truth; TF: transcription factor; G: gene.

To functionally assess GRN reconstruction, we examined the overlap in top-ranked regulons (i.e., regulator–target gene pairs) across methods (Figure 4C, S1C). Overall, regulon recovery was limited, with both regulator and target gene detection rates remaining low. Notably, a bias toward recovering effector (target) genes over transcriptional regulators was observed, likely reflecting the transcription factors’ lower and noisier expression levels in single-cell data.

We further benchmarked structural recovery using standard metrics from the BEELINE framework [3], including AUROC, AUPRC, and early precision-recall (EPR). Across all methods, overall accuracy and precision were low, especially in cyclic GRNs. SpliceJAC performed better on linear differentiation trajectories, showing higher AUROC and AUPRC scores than in cyclic settings (Figure 4D). In contrast, ScMomentum exhibited improved performance for cyclic dynamics, particularly in AUPRC and EPR metrics (Figure 4F). When restricting the analysis to activating edges only, the differences between linear and cyclic systems largely disappeared (Figure 4D–F).

Taken together, these results suggest that linear systems yield more precise network estimates, while cyclic dynamics remain more challenging to resolve. Nevertheless, across all conditions, the overall accuracy of weight estimation was suboptimal.

These findings indicate that while Jacobian-based methods can partially capture regulatory relationships, they struggle to reconstruct the GRN structure with high fidelity. The consistent bias toward target gene recovery and poor regulon overlap implies that additional methodological constraints or prior information may be necessary for improved inference. Given these limitations in structural recovery, it becomes essential to determine whether they stem from inherent methodological shortcomings or sensitivity to hyperparameter tuning. We, therefore, next assess the computational stability of these Jacobian inference methods.

### Jacobian Inference Sensitivity Results

An important aspect of method evaluation is assessing the sensitivity of Jacobian inference to key hyperparameters, including window size, step size, number of genes, and number of cells.

Window size showed minimal influence on the eigenvalue trends for the analytical method, ScMomentum, and Dynamo, which retained overall profile stability. In contrast, SpliceJAC displayed high variability, consistent with its direct reliance on local window-based calculations (Figure S2).

Step size exerted a more substantial influence on inferred dynamics than window size (Figure S2). For the analytical Jacobian, eigenvalue peak positions shifted with step size changes, and similar effects were observed for Dynamo and ScMomentum. However, larger step sizes led to reduced peak resolution in Dynamo. Correlation metrics exhibited a more complex, non-monotonic trend in response to step size, suggesting interactions between sampling granularity and underlying signal structure.

Gene number sensitivity was analyzed across acceptable ranges specific to each method. At low gene counts, global trends were preserved, though local variability emerged for SpliceJAC and ScMomentum (Figure S3A–C). Intermediate gene counts caused more pronounced effects with signal scaling, particularly in ScMomentum. For ScMomentum, shifts in transition peak features indicate the presence of an optimal gene count range for feature detection (Figure S3D–F).

Cell number influenced signal resolution across all methods, with higher cell counts generally improving trend definition (Figure S4). However, noise amplification accompanied signal gain, especially for Dynamo. These findings suggest a minimal threshold of cell numbers is required to achieve interpretable Jacobian estimates without introducing excessive noise.

For ScMomentum, which initializes Jacobian estimation through cluster-wise Hopfield networks, the number of clusters had a direct effect on eigenvalue profiles. Sensitivity analysis revealed stable behavior within specific clustering resolution ranges. We recommend practitioners test multiple clustering granularities to identify robust regions of stability.

Overall, our results indicate that while most Jacobian inference methods are broadly robust to hyperparameter variation, step size, and cell count have the most substantial impact on performance. These dependencies emphasize the importance of careful parameter tuning when applying Jacobian-based models to biological data.

In summary, our benchmarking analyses reveal that Jacobian-based methods differ considerably in their capacity to detect transitions, resolve oscillatory dynamics, and reconstruct regulatory structures. While some methods demonstrate robustness across data types and parameter settings, others exhibit high sensitivity to analysis conditions. These findings establish a foundation for guiding the application of Jacobian inference in single-cell dynamical modeling and underscore key considerations for future methodological refinement.

## Discussion

We evaluated both the phenomenological and structural integrity of Jacobians inferred from single-cell transcriptomic data. Our results show that ScMomentum and Dynamo effectively capture phenotypic transitions, as reflected in their eigenvalue spectra, while SpliceJAC is more sensitive to local sampling parameters. Application to real datasets further revealed that some methods can highlight dynamic cell states, although their ability to capture oscillatory behavior remains limited.

With respect to network reconstruction, we found that the structural accuracy of Jacobians remains low across methods. Regulon-level recovery is poor, and inferred weights often deviate significantly from ground truth distributions. Although the methods exhibit general robustness, our sensitivity analysis identifies key hyperparameters, such as cell number and gene selection strategy, that substantially influence performance and should be carefully considered during implementation. Additionally, since the methods are operational within small gene size ranges, it becomes more critical to prioritize sequencing throughput over depth. Nonetheless, features of the trends exist that are robust to even variable dataset size, given that they exceed a critical threshold.

A significant source of variation between methods lies in how genes are selected for inclusion. Splice-JAC relies on top variable genes filtered through scVelo, whereas Dynamo and ScMomentum allow explicit control over the genes used in Jacobian construction. Dynamo further enables users to specify regulators and effectors separately, typically derived from the principal components of the velocity field. In contrast, ScMomentum enforces a square Jacobian where regulators and effectors are identical. These design choices directly impact both interpretability and model flexibility.

The availability of RNA velocity has made Jacobian inference a feasible tool for exploring regulatory dynamics in single-cell data. However, significant architectural improvements are required to mitigate noise and improve reliability. Emerging methods aim to enhance RNA velocity estimation through Bayesian, deep learning, or dynamical modeling strategies [30, 31, 32, 33, 34, 35, 36, 37, 38, 39, 40]. These advances offer a foundation for generating more accurate Jacobians. Although this may lead to dense connectivity, regularization strategies can be incorporated to promote sparsity and interpretability [13, 41].

Additionally, integrating prior regulatory knowledge could further improve structural fidelity. In ecological and cellular systems, the use of known network topologies has proven effective [42]. For single-cell applications, GRN priors may be derived from curated databases or inferred through multiomic integration [43, 44, 45, 46, 47, 48]. Recent studies demonstrate the power of such approaches for sconstraining Jacobian inference and improving biological interpretability [49, 50, 51, 41].

In conclusion, we anticipate that Jacobian inference will become a key component in bridging single-cell transcriptomics with formal systems biology models. By enabling quantitative analysis of regulatory dynamics, these methods will support the development of mechanistic models of cell fate transitions and advance efforts in cell state control and therapeutic intervention.

## Resource availability

### Data and code availability

- **Data:** This study utilizes publicly available datasets, with access details provided in the Methods section. The Spermiogenesis dataset is available under accession number GSE221226, and the mouse pancreatic endocrinogenesis dataset is available under GSE132188.
- **Code:** all original code has been deposited in GitHub: https://github.com/balubao/Transition_Benchmark.git and is publicly available.
- **Additional information:** any additional information required to reanalyze the data reported in this paper is available from the lead contact upon request.

## Acknowledgements

We thank Robert Lehmann and Vincenzo Lagani for productive discussions and feedback. We acknowledge the KAUST Baseline Awards, with D.G.-C. supported by KAUST Baseline Award no. BAS/1/1093-01-01 and J.T. supported by KAUST Baseline Award no. BAS/1/1078-01-01. The graphical abstract was created with BioRender.com.

## Author Contributions

A.B. conceived the study. A.B. and J.P.B.T. conducted the experiments. R.K., H.A., and D.G.C. analyzed the data. N.K. and J.T. provided conceptual guidance. D.G.C. and J.T. secured funding and provided resources. All authors reviewed and approved the manuscript.

## Declaration of generative AI and AI-assisted technologies in the writing process

During the preparation of this work, the author(s) used ChatGPT to improve readability and flow. After using this tool/service, the author(s) reviewed and edited the content as needed and take(s) full responsibility for the content of the published article.

## Supplementary Figures

**Figure S1:**
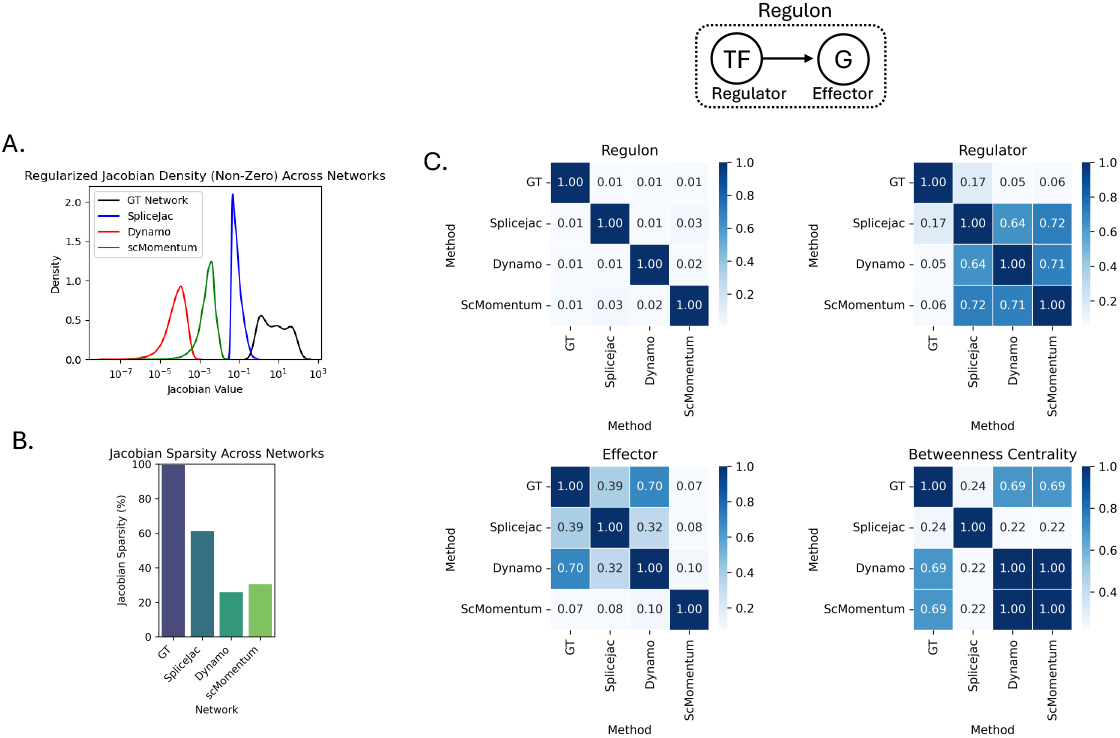
Structural Integrity (Cyclic): A. GRN weight distribution for the GT and Jacobians estimated by the other methods. B. Percentage of zeros in the Jacobians estimated between the different methods compared to the ground truth. C. The network substructure similarity was evaluated with respect to the top regulons (edge from regulator to effector), the Jaccard index is computed between the top effectors, regulators, and regulons in order of edge value (Top_N_regulons = 1000). The similarity in the top genes by betweeness centrality was also evaluated (Top_N_genes = 115). For Dynamo and scMomentum, the average Jacobian for all cells was used. All results correspond to the cyclic trajectory. GT: ground truth; TF: transcription factor; G: gene.

**Figure S2:**
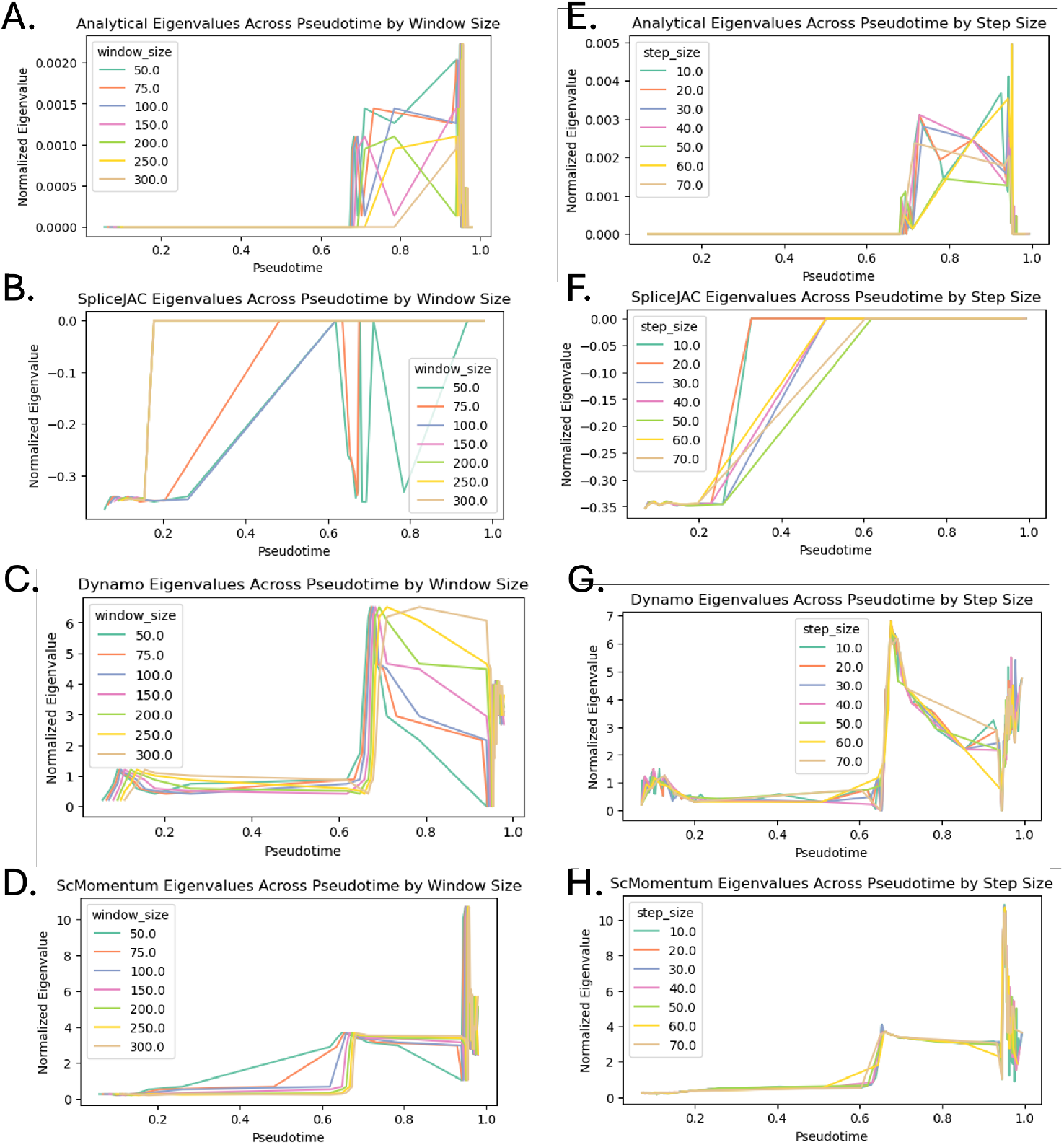
Time window parameter sensitivity: A. Panel shows the analytical jacobian leading eigenvalue trend along diffusion pseudotime for different window sizes as indicated by the figure legend. B-D. Same as A for the SpliceJAC, Dynamo, and ScMomentum jacobians. E-H. Same as A-D with different step sizes as indicated by the legends.

**Figure S3:**
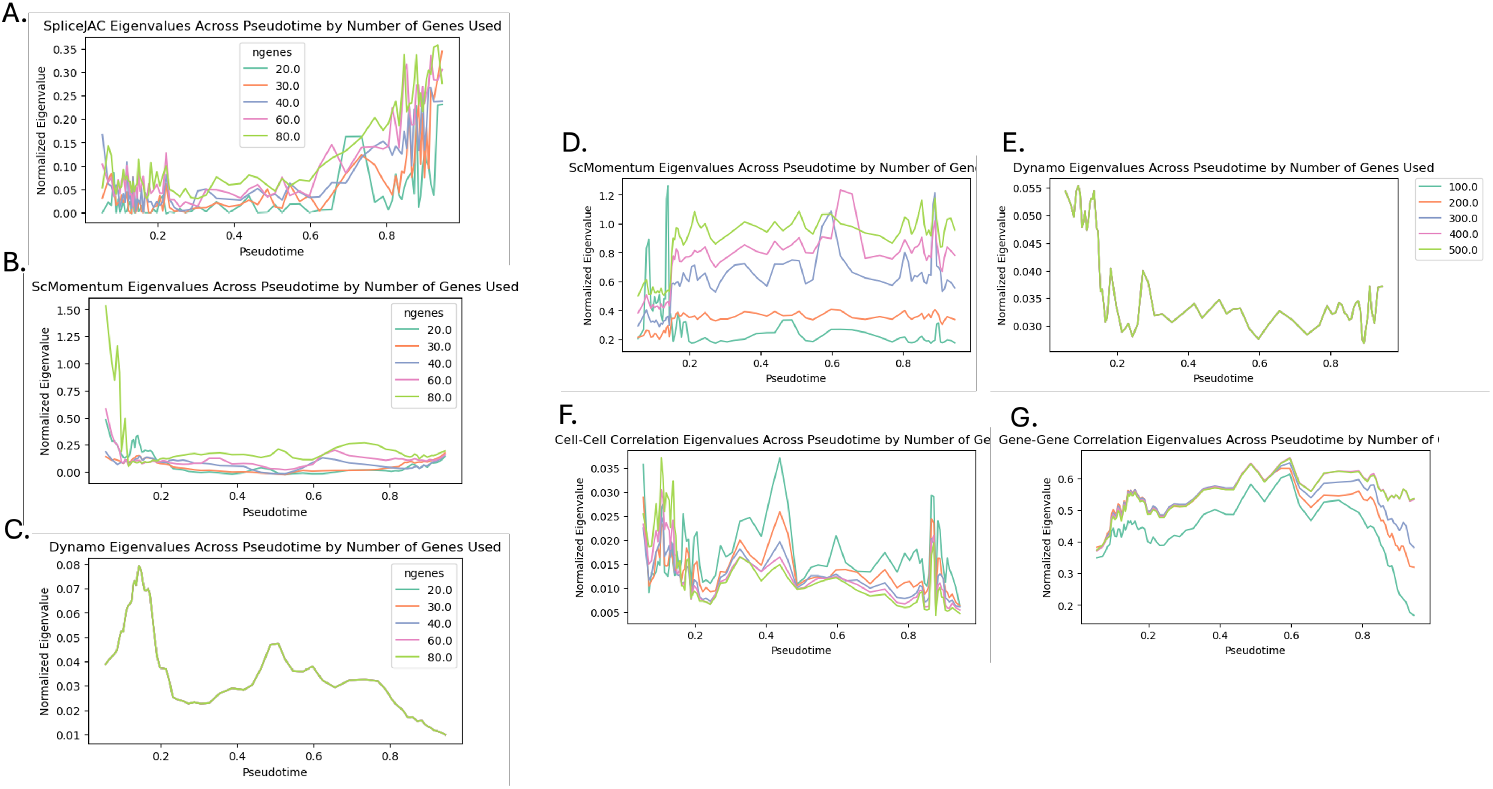
Number of Genes Sensitivity: A-C. Jacobian eigenvalue trend for low range of gene counts for (A) SpliceJAC, S(B) cMomentum, and (C) Dynamo. D-G. Metric trends for methods accepting intermediate number of genes, including jacobian leading eigenvalue trends from (D) scMomentum, (E) Dynamo, and average (F) cell-to-cell pearson correlation, and (G) gene-to-gene pearson correlation.

**Figure S4:**
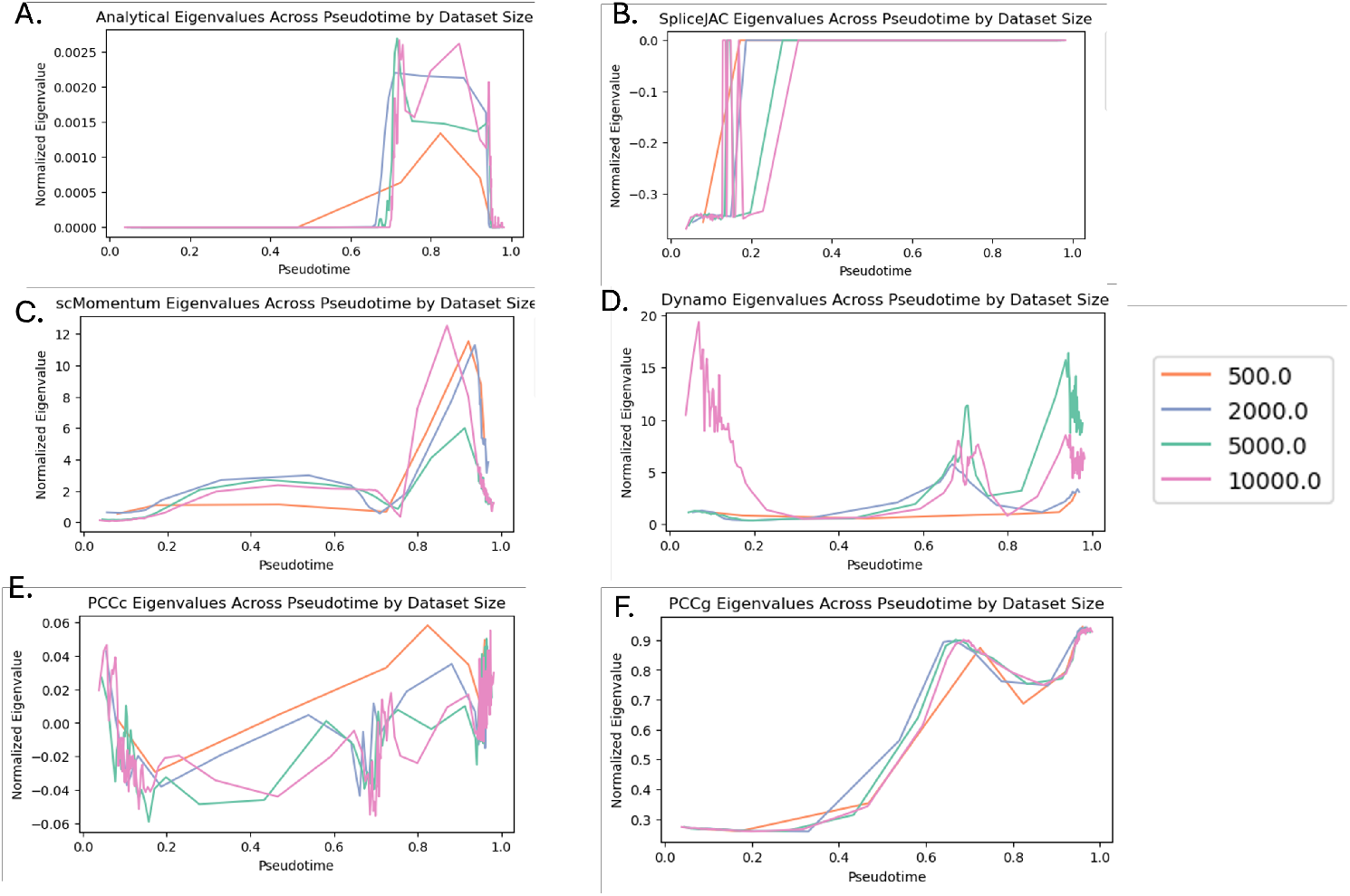
Number of cells Sensitivity: A. The trends for the analytical jacobian leading eigenvalue trend along diffusion pseudotime estimated using varying number of cells as indicated by the figure legend. B-D. Same as A for the jacobians estimated by SpliceJAC, ScMomentum, Dynamo. E-F. The average pearson correlation along pseudotime using different number of cells as indicated by the legend for (E) cell-to-cell correlation and (F) gene-to-gene correlation.

